# Heat-mediated manipulation of gene expression by IR-LEGO in the developing genitalia in *Drosophila*

**DOI:** 10.1101/2025.04.14.648785

**Authors:** Moe Onuma, Tatsuyuki Kumagai, Kentaro Hayashi, Yasuhiro Kamei, Aya Takahashi

## Abstract

Manipulating gene expression in a tissue-specific and temporally controlled manner is essential for understanding the function of the focal genes. Still, in many cases, the limited availability of specific promotors to drive ectopic manipulation remains a restricting factor in developing organs, even in *Drosophila*. Developing external genitalia is one such organ with a complex anatomical structure shaped by a joint regulatory network of many transcription factors. To overcome the restriction, we employed the infrared laser-evoked gene operator system (IR-LEGO), in which infrared laser (1,480 nm) irradiation induces gene expression under the control of a heat shock promoter. Pupal genital structures were irradiated at approximately 24 or 48 hours after puparium formation. We tested a range of laser power and depth to the target structure by a reporter assay using green fluorescent protein, which was induced under the control of the *heat shock protein 70* promoter (*hs*-*GAL4*). In previous studies, the IR-LEGO has been used as a tool to induce ectopic transgene expression. In this study, we attempted to knock down genes such as *yellow* (*y*) and *odd-paired* (*opa*) ectopically by RNAi using the GAL4/UAS system. The results demonstrated that this technique has a high potential in manipulating transcript abundance levels in small groups of cells in specific genital structures to unravel novel functions of genes involved in the morphogenesis of species-specific and rapidly evolving anatomical structures.

**Article summary:** Manipulating gene expression is essential for understanding the function of the genes at specific regions of the body. In particular, knowing how different genes contribute to shaping complex anatomical structures, such as insect external genitalia, is an intriguing question. While manipulating gene expression precisely at a specific time and space remains challenging, we show that we can knock down a gene in a specific group of cells in the developing genitalia of *Drosophila* pupae, using infrared laser irradiation. This is a new approach to investigating how activity of genes shapes the rapidly evolving male genitalia in insects.

## Introduction

Manipulating gene expression is essential for understanding the functions of the focal genes. In particular, regulating gene expression in a tissue-specific and temporally controlled manner is often required to overcome complexity caused by pleiotropic effects from multiple expression domains of the focal genes. Various genetic tools have been developed to achieve this objective. A groundbreaking technique, the GAL4/UAS system utilizing yeast galactose-inducible system has proven to be versatile and highly effective in manipulating transcription activation in *Drosophila melanogaster* (Brand and Perrimon 1993) and has been widely used. It can also be used in a slightly limited manner in mice and zebrafish (Asakawa and Kawakami 2008; Halpern et al. 2008). However, despite its widespread application, finding the correct GAL4 driver for a specific purpose is not trivial when the drivers that are activated at a specific time and space of interest are not available due to pleiotropy. Further methodological refinement is necessary to conduct precise spatial and temporal manipulation of the gene expression levels in such cases.

External genitalia, especially that of males, is one of the most complicated and rapidly diversifying anatomical structures among animal body parts (Eberhard 1985; Hosken and Stockley 2004; Simmons 2014). Recent studies using pairs of closely related *Drosophila* species have highlighted the expression diversifications of genes that are responsible for interspecific morphological differences in specific genital structures (Nagy et al. 2018; Hagen et al. 2019; Ridgway et al. 2024). However, the molecular basis of the unique morphological evolution of genital structures remains largely elusive.

The whole terminalia (genitalia and analia) develops from the genital imaginal disc, and a higher-order morphogenesis takes place throughout pupal development. While genetic regulation in the imaginal discs has been described in more detail (Gorfinkiel et al. 1999; Keisman and Baker 2001; Chatterjee et al. 2011; Ridgway et al. 2024), a high throughput in situ hybridization assay of the pupal terminalia (including genital and anal structures) of male *D. melanogaster* has depicted the expression landscapes of over 100 transcription factors (Vincent and Rice et al. 2019). Also, the morphogenesis anatomy of male pupal terminalia in 12 *Drosophila* species with diverse morphology has been described and compared in detail (Urum et al. 2024; Urum and Preger-Ben Noon 2025). The functional assays of the transcription factors and other genes would apparently be the next step in tackling the question of how gene regulatory networks shape complex morphogenesis and facilitate diversification. Such an approach requires precise manipulation of gene expression at specific substructures or cell populations at specific timings. However, with currently available genetic tools, manipulating gene expression *in vivo* in certain visible domains of developing genitalia is still challenging, especially for the transcription factors with pleiotropic roles. Motivated by this demand, we have developed a heat-mediated manipulation system of gene expression in the developing genitalia of *D. melanogaster*.

In this study, we employed the infrared laser-evoked gene operator system (IR-LEGO, Kamei et al., 2009), in which infrared laser light irradiation induces gene expression under the control of a heat shock promoter. The method has been successfully applied to *Caenorhabditis elegans* (Kamei et al. 2009; Suzuki et al. 2013; Suzuki et al. 2022), *Danio rerio* (Deguchi et al. 2009; Kimura et al. 2013), *Oryzias latipes* (Deguchi et al. 2009; Kobayashi et al. 2013; Okuyama et al. 2013; Shimada et al. 2013), *Xenopus laevis* (Kawasumi-Kita et al. 2015; Hasugata et al. 2018), *Pleurodeles waltl* (Kawasumi-Kita et al. 2015), *Arabidopsis thaliana* (Deguchi et al. 2009; Hwang et al. 2019; Tomoi et al. 2023), *Physcomitrium patens* (Tomoi et al. 2024), *Daphnia magna* (Shimizu et al. 2024), and *Drosophila melanogaster* (Miao and Hayashi 2014). In this study, we combined the inducible RNAi knockdown system with IR-LEGO for the first time.

In brief, the pupal terminalia at approximately 24 or 48 hours after puparium formation (h APF) was irradiated by an infrared laser. The efficiency of the heat shock induction was examined using *hs-GAL4*/*UAS-GFP* flies, in which the green fluorescent protein (GFP) is induced under the control of the *heat shock protein 70* promoter. Furthermore, we knocked down the *y* and *opa* genes by the inducible RNAi system. We demonstrate that the technique is promising in manipulating transcript abundance in small groups of cells in specific genital structures and can be employed to unravel novel functions of genes in the morphogenesis of rapidly evolving genital structures.

## Materials and Methods

### Fly Stocks

The following *D. melanogaster* strains were used in the present study: *hs-GAL4* (#106-509) from Kyoto *Drosophila* Stock Center, *UAS-CD4-tdGFP* (#35838) from Bloomington *Drosophila* Stock Center, *UAS-y*-*RNAi* (3757R-1) and *UAS-opa*-*RNAi* (1133R-3) from National Institute of Genetics. The NIG-Fly assessment of phenotypes induced by *Act5C*-*GAL4* at 28°C showed “yellow body color” and “lethal” for 3757R-1 and 1133R-3, respectively, indicating sufficient efficiency of RNAi knock down. All flies were maintained under the 12 h: 12 h light-dark condition at 25 ± 1°C on the standard corn medium.

### Sample preparation

Male and female pupae were collected at the white pupal stage by sorting gonad size and placed in a humid chamber at 25 °C. The puparium at the abdominal tip was removed by fine forceps at 24 ± 1 h APF or 48 ± 1 h APF and placed onto a glass bottom petri dish (35 mm dish diameter, 14 mm glass diameter, glass thickness 0.16−0.19 mm, D11130H, Matsunami, Japan) through a hole made by 1,000 uL pipetman tip to a 250−300 μL drop of solidified 1.5% agarose (Figure 1). The hole needs to be tilted to allow light penetration through the tissues to visualize the target structures. Distilled water was supplied to fill the space between the specimen and the glass.

**Figure 1.**
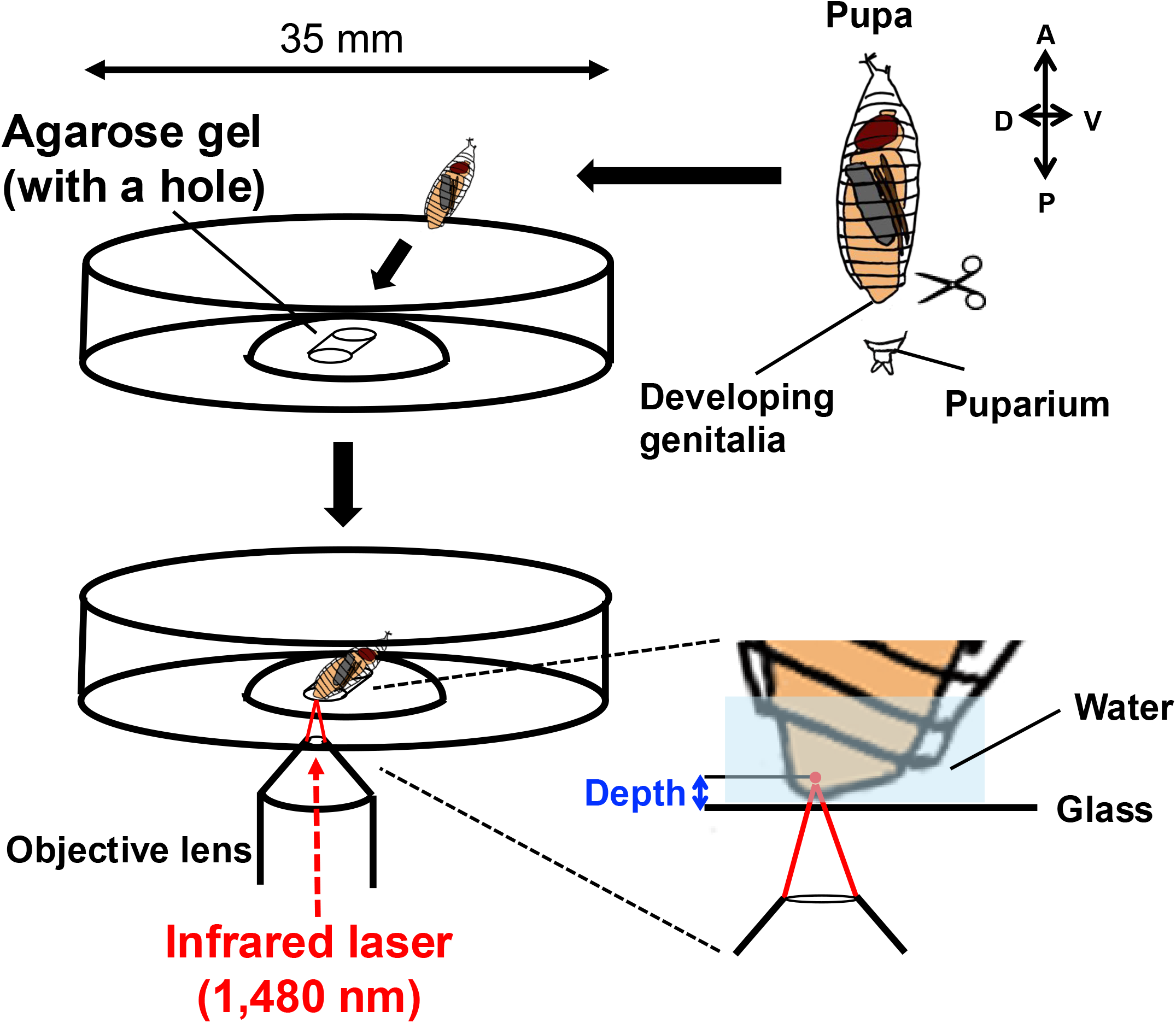
Sample preparation and laser irradiation of the pupal terminalia. Puparium at the abdominal tip was removed to expose the developing genitalia. The pupa was placed onto a glass bottom petri dish through an angled hole made to a drop of solidified agarose. Infrared laser (1,480 nm) was applied through the objective lens and irradiated the target cells at a certain distance (depth) from the glass.

### IR-LEGO system

An inverted microscope (IX-81, Olympus, Japan) was equipped with an IR-LEGO optical unit (custom assembled, Sigma-Koki, Japan). An objective lens, UApo340 20X (NA = 0.75 UV) (custom made, Olympus, Japan), was used to visualize the target tissues and focus irradiation with an infrared laser (1,480 nm). The distance between the bottom glass (marked by an ink pen) and the focal cells (Figure 1) was measured using the images captured by a CCD camera (ORCA-Flash 4.0, C11440, Hamamatsu Photonics, Japan) attached to the inverted microscope. Metamorph Software (Molecular Devices, Japan) was used to measure the distance (depth). The laser power (dBm) was calibrated by using a Head Sensor for Laser Power Meter 10A (Ophir 7Z02637, Japan) and a Vega Laser Power & Energy Meter (Ophir 7Z01560, Japan). A 6−8 mW laser through the objective lens was emitted to heat the live specimen for 60 s.

### Observation of fluorescent signal and morphology after irradiation

Irradiated pupa was kept in the agarose gel inside the petri dish with the lid closed to avoid desiccation. The petri dish was placed at 25°C until eclosion. For the *hs*-*GAL4*/*UAS-CD4-tdGFP* individuals, the fluorescent signal was observed on the following day by a band path filter set (470−495 nm excitation filter, 510−550 nm emission filter, 505 nm dichroic mirror) equipped with an inverted microscope (IX-81, Evident, Japan). Fluorescent images were taken with a CCD camera (ORCA-Flash 4.0, C11440, Hamamatsu Photonics, Japan) attached to the microscope.

Adults that emerged from the pupae after irradiation were taken out of the petri dish and placed into 70% ethanol at 24−48 hours after eclosion. The *hs*-*GAL4*/*UAS*-*y*-*RNAi* and the *hs*-*GAL4*/*UAS*-*opa*-*RNAi* pupae were dissected in 70% ethanol, and the periphallic genital organs of males or the hypogyniums of females were mounted in 50% Hoyer’s solution (Hoyer’s medium: acetic acid = 1:1) and incubated at 60°C overnight. Images of genital organs were captured by a CCD camera (DP73, Evident, Japan) attached to an inverted microscope (IX73, Evident, Japan).

### Linearity scores of the proximal bristle alignment on the surstylus

The linearity of the five most proximal bristles on the surstylus was evaluated by the proportion of variance explained by the first principal component extracted by the principal component analysis (PC1). The bases of the bristles surrounded by a socket cell on the photo images were manually marked, and their X and Y coordinates were obtained using Fiji (Schindelin et al. 2012). The PC1 of the coordinates represents a line that minimizes perpendicular distances from all points. The proportion of the variance explained by this component is an indicator of the distances from the line. Therefore, bristle coordinates with a score closer to 1 are more linearly aligned.

## Results

### Optimal IR-LEGO conditions for developing pupal genitalia in *Drosophila*

The optimal condition to induce heat-shock mediated gene expression varies among organisms and possibly among genetic systems employed (Deguchi et al. 2009). The laser power, irradiation duration, and the distance between the coverglass and the focal cells (hereafter depth) are the factors affecting the efficiency because of the attenuation of laser power due to the absorption of water in the tissue or gel in front of the focal cells. Previous investigations have shown that longer irradiation (i.e., 60 s) results in a more stable heat shock response compared to shorter durations, and a higher laser power will have a higher risk of damaging the cell (Kamei et al. 2009; Tomoi et al. 2024). Thus, for the developing pupal genitalia, we tested 6, 7, and 8 mW laser power with a duration of 60 s, targeting cells at various depths.

To find the optimal condition, the pupal genitalia of *hs-GAL4*/*UAS-CD4-tdGFP* at 48 ± 1 h APF, when structures of male and female external genitalia such as surstylus (clasper) and hypogynial valve (ovipositor) become visible (Glassford et al. 2015; Smith et al. 2020; Urum et al. 2024), were targeted. The laser was applied to two well-focused positions (chosen arbitrarily) on the right side of the surstylus (for a male, Figure 2A) or the hypogynial valve (for a female, Figure 2B), leaving the other side intact. The GFP signal was visible the next day when the heat shock-mediated GFP induction was successful (Figure 2BB’DD’).

**Figure 2.**
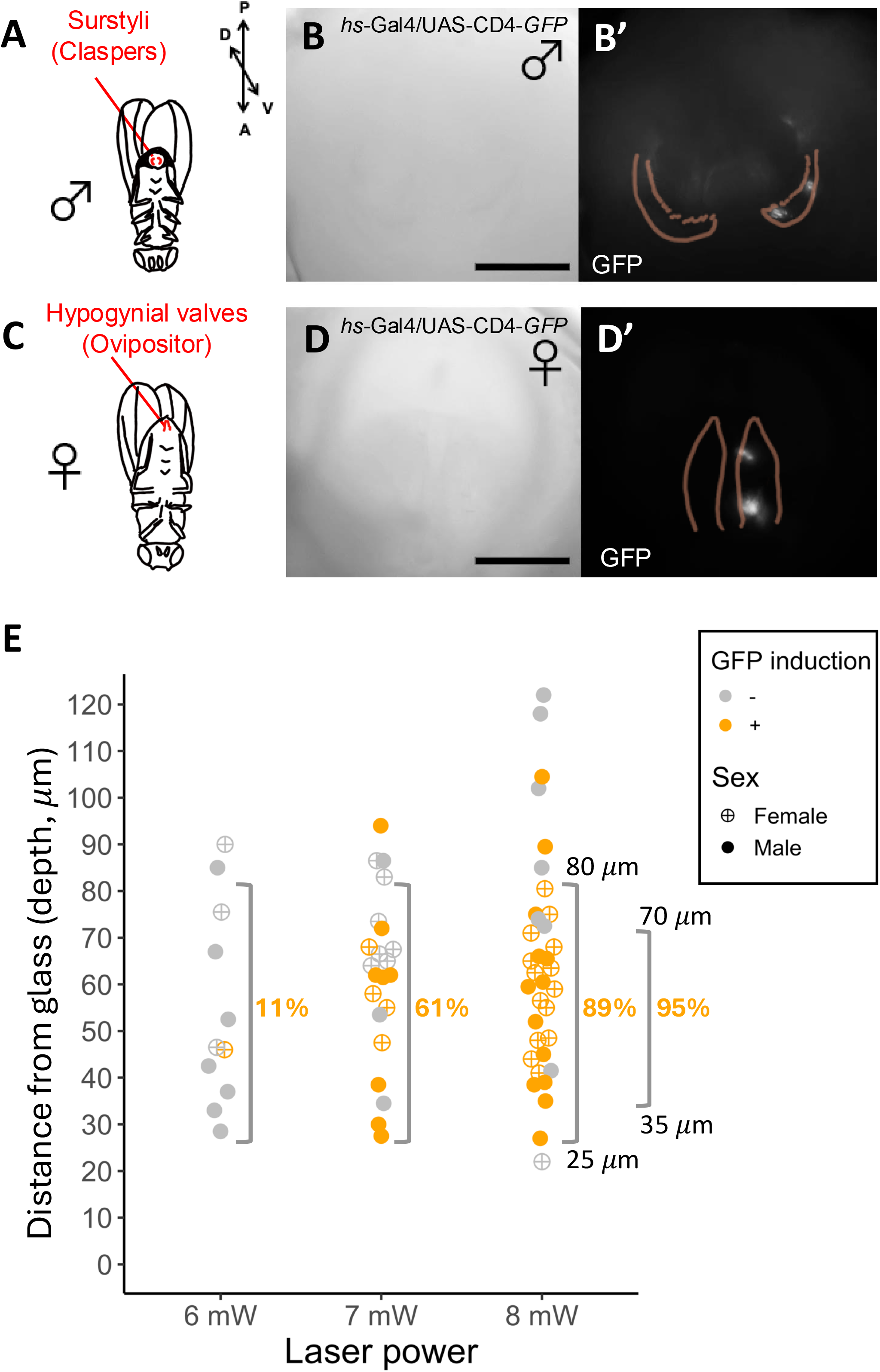
Testing experimental conditions for IR-LEGO in *Drosophila* pupae. (A) Surstyli of an adult male. (B) Bright field image and (B’) GFP signal of the pupal genitalia one day after infrared laser irradiation to the surstylus of a male *hs-GAL4/UAS-CD4-tdGFP*. Outlines of the pair of sursyli are indicated in B’. (C) Hypogynial valves in an adult female. (D) Bright field image and (D’) GFP signal of the pupal genitalia one day after infrared laser irradiation to the hypogynial valve of a female *hs-GAL4/UAS-CD4-tdGFP*. Outlines of the pair of valves are indicated in D’. (E) Presence and absence of GFP signal one day after irradiation by a 6, 7, and 8-mW laser in the target cells at various distances (depths) from the glass. Percentages of GFP-positive samples are shown within the ranges indicated by brackets. Scale bars indicate 100 μm.

The depth of the target structure could not be precisely controlled due to the variation in the shape of the puparium. Therefore, we tested the responses of the target groups of cells at various depths using different laser power settings. The results are shown in Figure 2E and Supplementary Table S1. The minimum depth, which was restricted by the pupal cuticle covering the developing structures, among the samples was 22 *μ*m. The samples with at least one detectable signal were scored as GFP-positive. The results indicated that GFP signal was detected in 89% of the samples within the dept range 25−80 *μ*m when irradiated using 8 mW infrared laser beam for 60 s. The success rate increased to 95% when the depth range was confined to 35−70 *μ*m. The eclosion rate of the irradiated samples was 91.9% (*N* = 74). According to these results, the samples with depth > 80 *μ*m were discarded in the following experiments.

### The heat-mediated RNAi knock down of the *y* gene in small groups of cells

Since the RNAi knock down by IR-LEGO has not been conducted before, we tested the effect of heat-mediated knock down using the *y* gene, whose classic mutants exhibit yellow body color due to the lack of an ability to synthesize dark-colored pigments (Figure 3CH). We used the same GAL4/UAS system as in the case of the GFP reporter assay (Figure 2), and the *hs-GAL4*/*UAS-y-RNAi* individuals were subjected to infrared laser irradiation targeting one side of the paired surstyli (Figure 3AB) or hypogynial valves (Figure 3EF). Three positions close to the tip of those structures at 48 ± 1 h APF (Figure 3BF) were irradiated by an 8-mW infrared laser beam for 60 s (Supplementary Table S2). After eclosion, a small number of yellow-colored bristles, like those of *y*^1^, were observed only on the irradiated sides of those structures (Figure 3DD’HH’). This phenotype was not observed in male and female individuals in the GFP reporter assay subjected to the same irradiation treatment and showed a clear GFP signal (Supplementary Figures 1 and 2). These results indicated that infrared light irradiation can be used to induce RNAi knockdown by the heat-mediated inverted repeat RNA transcription.

**Figure 3.**
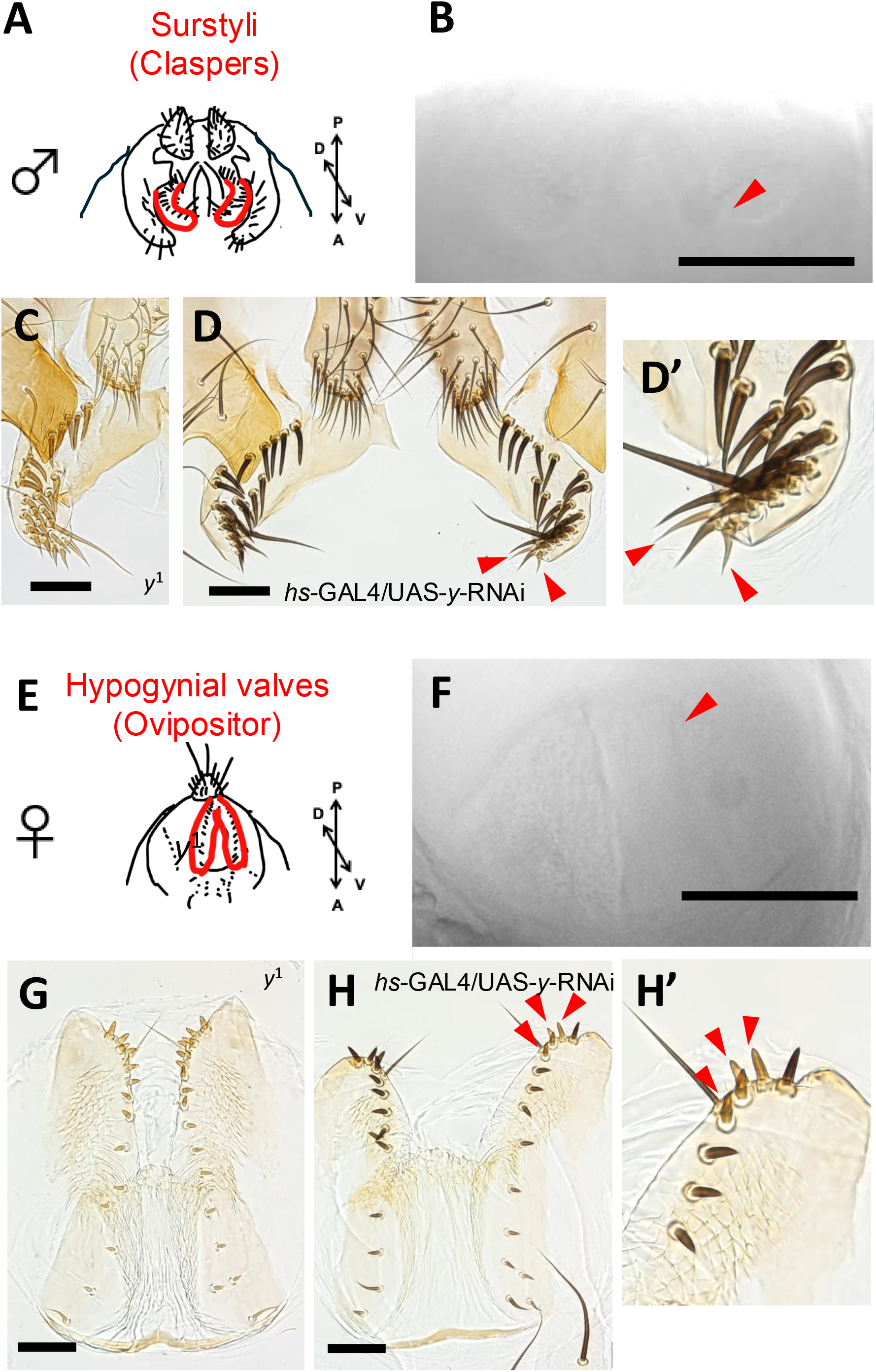
RNAi knock down of *y* by IR-LEGO in pupal genitalia. (A) Surstyli of an adult male. (B) Bright field image of the irradiated surstylus at 48 ± 1 h APF. Arrowhead indicates the approximate location of irradiated positions (three times) by an 8-mW laser for 60 s. (C) Surstyli of *y*^1^ mutant. (D) Left (intact) and right (irradiated) surstyli of an *hs-GAL4*/*UAS-y-RNAi* male after eclosion. (D’) Magnified image of the right surstylus. Arrowheads indicate bristles with reduced black pigmentation. (E) Hypogynial valves of an adult female. (F) Bright field image of the irradiated valve at 48 ± 1 h APF. Arrowhead indicates the approximate location of the irradiated positions (three times) by an 8-mW laser for 60 s. (C) Hypogynial valves of *y*^1^ mutant. (D) Left (intact) and right (irradiated) valves of an *hs-GAL4*/*UAS-y-RNAi* female after eclosion. (D’) Magnified image of the right valve. Arrowheads indicate bristles with reduced black pigmentation. Scale bars indicate 100 *μ*m.

### The heat-mediated RNAi knock down of the *opa* gene at the border region of epandrial ventral lobe and surstylus

Motivated by the accumulating information on transcription factor expression domains in developing fly terminalia (Vincent and Rice et al 2019), we aimed to analyze a transcription factor gene, *opa*, by knocking it down using the IR-LEGO system. This gene is expressed exclusively in the surstylus at 48 h APF and in the medial portion of the epandrial ventral lobe / surstylus domain (a single continuous epithelium) at 28 h APF (Figure 4A), which suggests that it may play a role in identifying presumptive surstylus tissue prior to its cleavage from the epandrial ventral lobe (Vincent and Rice et al. 2019). The physical separation of epandrial ventral lobe and surstylus begins at around 28 h APF (Urum et al. 2024).

**Figure 4.**
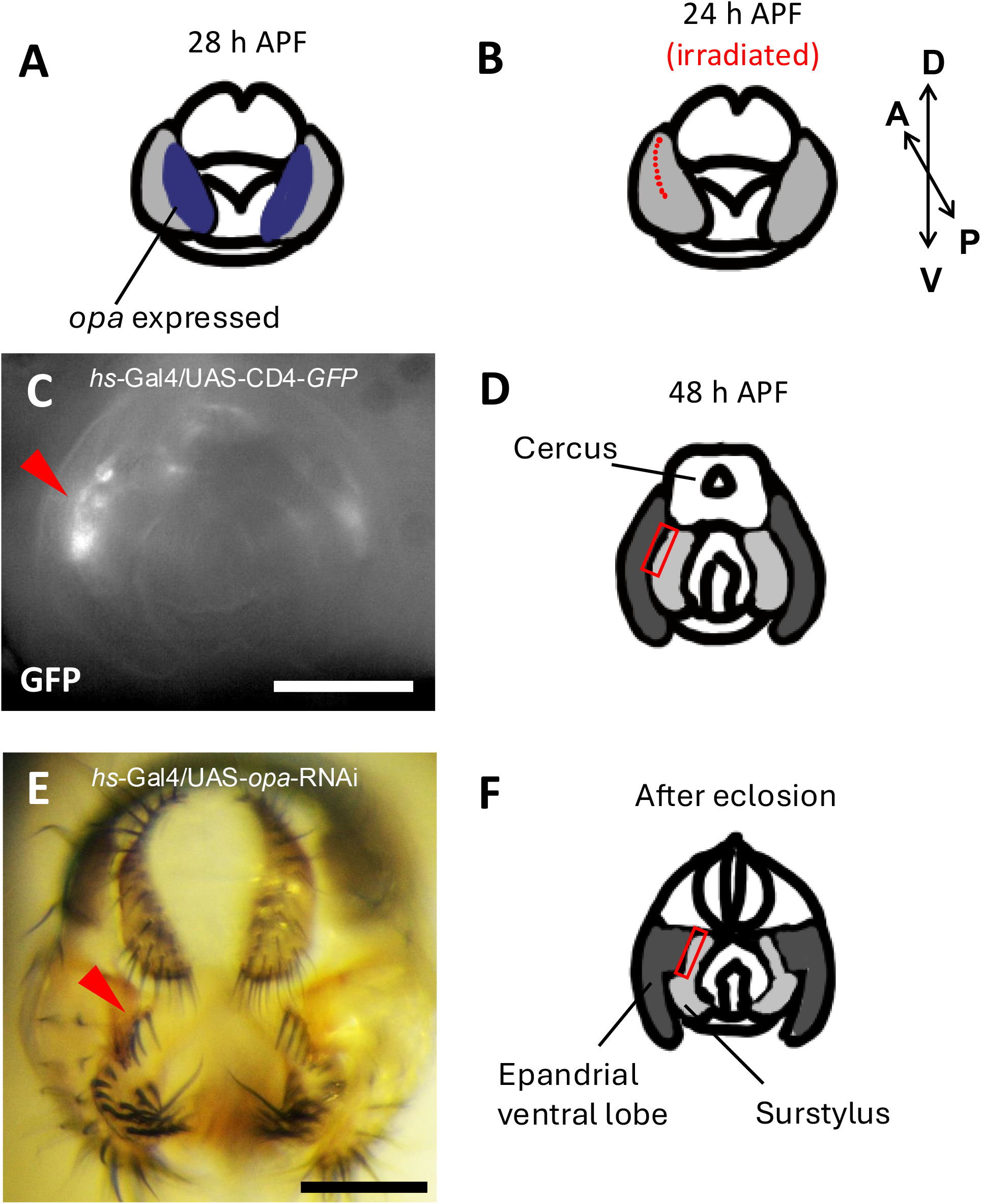
RNAi knock down of *opa* by IR-LEGO in pupal surstylus. (A) Schematic image of the pupal terminalia at 28 h APF with *opa* expression domains (indicated in blue) in accordance with Vincent et al. (2019). (B) Schematic image of the pupal genitalia at 24 h APF. Light grey indicates joint epandrial ventral lobe / surstylus primordia. Red dots indicate approximate positions irradiated by an 8-mW laser for 60 s (ten times). Cleavage of the epandrial ventral lobe and surstylus initiates around this developmental time point. (C) GFP signal of the pupal genitalia one day after infrared laser irradiation at positions shown in B of *hs-GAL4/UAS-CD4-tdGFP*. Arrowhead indicates the affected area. (D) Schematic image of pupal genitalia at 48 h APF. Dark grey indicates epandrial ventral lobe and light grey indicates surstylus. Red rectangle indicates the affected area inferred from C. (E) Adult male terminalia of *hs-GAL4*/*UAS-opa-RNAi* eclosed after irradiation at positions shown in B. Arrowhead indicates the affected bristle alignment. (F) Schematic image of adult genitalia after eclosion. Dark grey indicates epandrial ventral lobe and light grey indicates surstylus. Red rectangle indicates the affected area inferred from E. Scale bars indicate 100 *μ*m.

To investigate the predicted role of *opa*, it is essential to manipulate the transcript levels at the boundary region. However, approaches using regulatory sequences require elaborate genetic engineering to confine specific targets for manipulation. Instead, we employed IR-LEGO and conducted the *opa* gene knock down targeting the epandrial ventral lobe / surstylus boundary, the epithelial area close to the future cleavage site, at 24 ± 1 h APF. The 8-mW laser was applied to 10 positions for 60 s on the right or left side of the surstylus (Figure 4B). The side with a clearer focus and a smaller depth was chosen.

The response to heat shock at this timing and location was confirmed by the GFP reporter (Figure 4CD). In adult flies after eclosion, we found that the alignment of the proximal bristles on the surstylus was slightly disrupted on the irradiated side (Figure 4EF). To quantify this subtle change, the linearity scores of the rows of the five most proximal bristles on both left and right sides were obtained after dissecting the adult fly and mounting the periphallic organs on slides, as in Figure 5A. An example of the marked positions of the bristles and the linearity scores is shown in Figure 5BC.

**Figure 5.**
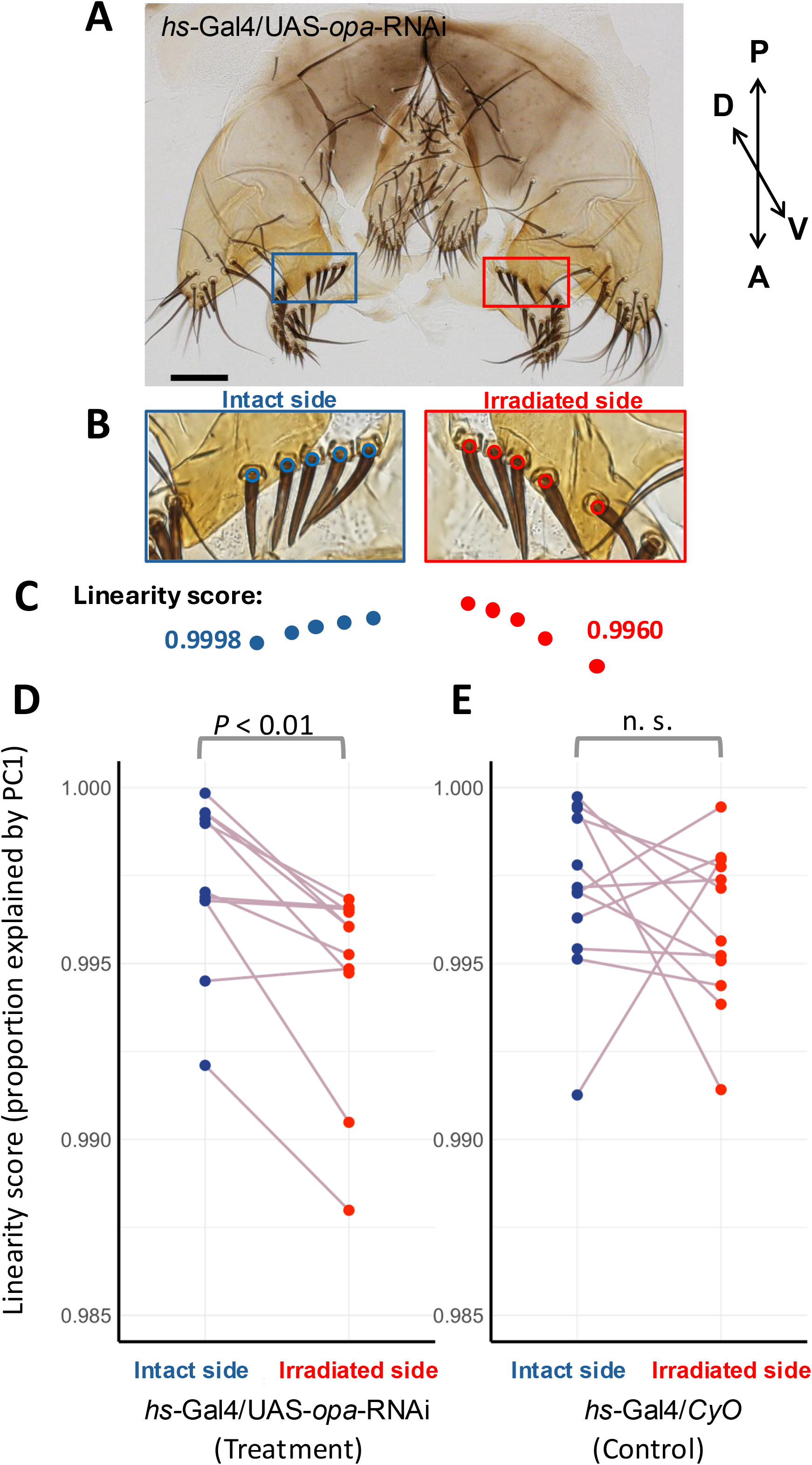
Quantification of the effect of *opa* knock down on surstylus bristle alignment. (A) A sample of the periphallic genital organs of *hs-GAL4*/*UAS-opa-RNAi* subjected to the IR-LEGO treatment shown in Figure 4. Phallic organs were removed and mounted on a slide. Red and blue rectangles indicate most proximal bristle rows on irradiated and intact sides, respectively. Scale bars indicate 100 *μ*m. (B) Marked positions of the bases of bristles surrounded by a socket cell. Red and blue circles represent bristle positions of irradiated and intact sides. (C) Alignments and linearity scores of the marked bristle positions of the irradiated (red) and intact (blue) sides in B. (D) Linearity scores (proportions of variance explained by PC1 of the bristle position coordinates) of the irradiated and intact sides of the *hs-GAL4*/*UAS-opa-RNAi* males subjected to the IR-LEGO treatment shown in Figure 4. (E) Linearity scores of the irradiated and intact sides of *hs-GAL4*/*CyO* control males subjected to the same treatment as in D.

The linearity scores were lower in the irradiated side compared to the intact side within the *hs-GAL4*/*UAS-opa-RNAi* individuals (Figure 5D, paired Wilcoxon signed-rank test, p < 0.01, Supplementary Table S3), indicating that the *opa* knock down at the presumptive boundary region between the epandrial ventral lobe and surstylus perturbs the surstylus bristle alignment close to the boundary in an adult fly. To confirm that the heat damage is not the cause of the perturbation, we investigated the *hs-GAL4*/*CyO* control individuals, in which there was no significant difference between the two sides (Figure 5E).

## Discussion

Our experiment succeeded to employ the IR-LEGO system for knocking down a gene at a specific timing and space in the developing genitalia of *Drosophila*. In our procedure, the target spot is irradiated for 60 s with an 8-mW infrared laser, while the previous study targeting the migrating tracheal cells of *Drosophila* embryos used a 44-mW laser for 1 s (Miao and Hayashi 2014). Many other previous studies have also employed 0.5−1-s irradiation (Deguchi et al. 2009; Kimura et al. 2013; Kawasumi-Kita et al. 2015) but longer durations have been applied to live cells of plants as they are immobile (Hwang et al. 2019; Tomoi et al. 2023; Tomoi et al. 2024). After checking the immobility of the pupal genital structures at APF 24 and 48 h upon infrared laser irradiation, we applied a longer irradiation duration (60 s) with lower laser power (6−8 mW), which was suggested in previous studies (Kamei et al. 2009; Tomoi et al. 2024) to ensure a more robust induction of heat-mediated transcription. However, pupae at later stages become mobile inside the puparia, therefore, irradiation condition may need to be adjusted.

Pupal epidermis is covered by a thin transparent pupal cuticle and the genital primordia develops near the posterior end of the cuticle. Thus, developing genitalia has been a favorable material to conduct *in vivo* imaging through the lens placed outside the cuticle (Suzanne et al. 2010; Kuranaga et al. 2011; Sato et al. 2015). With the IR-LEGO system, the cells in the focal plane can be irradiated *in vivo* through water and aquatic tissues inside the cuticle. Since the genital tissues are not completely transparent, the cells close to the surface are the most effective target. Although IR-LEGO has a resolution of targeting single cells, those in the distant position from the surface will be difficult to irradiate with sufficient resolution. We aimed to focus on the cells at or close to the surface. The transcription induction in our system is confined to small groups (likely to be < 5) of cells (Figure 2B’D’). For targeting a larger epithelial area, irradiating multiple sites is necessary as in a case of knocking down *opa* along the epandrial ventral lobe / surstylus boundary (Figure 4A).

The phenotypic effect of the heat-mediated manipulation of gene expression can be assessed with high sensitivity by comparing left and right sides when one side is irradiated leaving the other side intact. The heat shock response is most strongly induced when the cell is experiencing severe heat stress (Kamei et al. 2009). Therefore, it is necessary to be aware that some cells can be damaged by heat or affected by the general heat shock response of the cell. In embryos both with and without *hs*-*branchless*, inhibitory effect on terminal branch formation during the dorsal tracheal development was observed, indicating that the effect was due to irradiation itself (Miao et al. 2014). Likewise, by irradiating samples with and without RNAi-inducing transgene would be effective in controlling for the potential effect by heat shock response. In our study, we dissected the adults eclosed from heat-mediated induction of GFP as a control for *y* knock down (Supplementary Figures 1 and 2) and used *CyO* control in the case of *opa* knock down experiment (Figure 5D). These controls are essential for the IR-LEGO experiments.

The GAL4/UAS system has been widely used to manipulate gene expression at specific timing and space when an appropriate GAL4 driver is available. *hs*-*GAL4* has also been a useful tool to induce ectopic expression or RNAi knock down by combining it with various transgenes with the UAS promotor (Bainton et al. 2005; Kim et al. 2018; Seong et al. 2019). Inducing heat shock ubiquitously at the whole organism level is possible by typically placing late-stage pupae or adults at 37°C for 20−60 min (e.g., Armstrong et al. 2006; Nakayama et al. 2014). In this study, we successfully showed that the IR-LEGO heat-mediated manipulation via *hs*-*GAL4* can be performed at a confined position and timing during the genitalia morphogenesis by the *y* RNAi. Compared to other genes such as *ebony* and *tan* involved in cuticle pigmentation, *y* is known to be expressed at earlier timing during the pupal period. In the pupal wings, the transcript is most abundant at around 52 h APF (Sobala and Adler 2016) and declines while the protein becomes abundant (Riedel et al. 2011; Hinaux et al. 2018). In the abdomen and thorax, Yellow protein was detected at around 60−80 h APF and expression in cells associated with bristles have been also reported (Wittkop et al. 2002; Hinaux et al. 2018). Transcription of inverted repeat sequence of this gene at 48 h APF in our study was effective in reducing the dark pigmentation of the bristles on female and male genital organs, which coincide with these previous studies.

Among many transcription factors expressed during genital morphogenesis, *opa* is a transcription factor that is presumed to be involved in shaping the boundary between epandrial ventral lobe and surstylus (Vincent and Rice et al. 2019). These two structures develop from shared primordia and start separating by 28 h APF (Urum et al. 2024). *opa* marks the surstylus tissue at this time point and retains its surstylus specific expression up to 48 h APF at least (Vincent and Rice et al. 2019). We have irradiated the boundary epithelial tissues at 24 h APF, before the cleavage begins and found that the knock down of *opa* at this timing and location affects the proximal bristle alignment of the surstylus (Figures 4 and 5). The proximal half of the surstylus is attached to the epandrial ventral lobe and the proximal bristles are located along the edge of the medial surface of the surstylus close to the protruded epandrial posterior lobe. The knock down of *opa* in cells close to the boundary may disrupt the fate of some presumptive surstylus cells and perturbs the linearity of the bristle alignment on the surstylus. There were no apparent differences in the shape or size of the posterior lobes between the irradiated and intact sides, suggesting the possibility that *opa* is primarily determing the fate of the surstylus cells and not involved in the epandrium morphogenesis to a large extent. Further experiments to induce knock down or ectopic expression of this gene at different cells and time using this system would elucidate its role in genital morphogenesis.

In summary, we showed the potential of using IR-LEGO system to induce heat-mediated transcription of inverted repeat sequences through the GAL4/UAS system to conduct RNAi knockdown. While many studies focus on imaginal discs, boundary formation and tissue differentiation continue to take place through the pupal period to form complex genital structures that are unique to each species. Manipulating gene expression at the focal position is certainly an attractive method without relying on the availability of transgenic strains with regulatory sequences that can target specific tissues at specific timing. Furthermore, advanced genetic tools that incorporate the heat shock promotor, such as *hs-Cre* (i.e., Kobayashi et al. 2013), *hs-FLP*, and *hs-Cas9*, can be tested immediately using this system.

## Data availability

All data are provided in the Supplemental Tables S1−5. Raw images supporting these analyses are provided as Supplementary_files available at FigShare.

## Acknowledgments

We thank Misako Saida at National Institute of Basic Biology for her technical assistance in using the IR-LEGO facility.

## Funding

This work was supported by NIBB Collaborative Research Program (24NIBB515) and KAKENHI 23K27221 and 24K21994 to AT, KAKENHI 20H05891 and 23K26510 to YK, and KAKENHI 24KJ0181 to MO.

